# Beyond the first glance: How human presence enhances visual entropy and promotes spatial learning

**DOI:** 10.1101/2025.05.29.656764

**Authors:** Tracy Sánchez Pacheco, Debora Nolte, Sabine U. König, Gordon Pipa, Peter König

**Affiliations:** Institute of Cognitive Science, University of Osnabrück, Wachsbleiche 27, 49090 Osnabrück, Germany; Department of Neurophysiology and Pathophysiology, University Medical Center Hamburg-Eppendorf, Martinistr. 52, 20246 Hamburg, Germany

## Abstract

Spatial learning emerges not only from static environmental cues but also from the social and semantic context embedded in our surroundings. This study investigates how human agents influence visual exploration and spatial knowledge acquisition in a controlled Virtual Reality (VR) environment, focusing on the role of contextual congruency. Participants freely explored a 1 km^2^ virtual city while their eye movements were recorded. Agents were visually identical across conditions but placed in locations that were either congruent, incongruent, or neutral with respect to the surrounding environment. Using Bayesian hierarchical modeling, we found that incongruent agents elicited longer fixations and significantly higher gaze transition entropy (GTE), a measure of scanning variability. Crucially, GTE emerged as the strongest predictor of spatial recall accuracy. These findings suggest that human-contextual incongruence promotes more flexible and distributed visual exploration, thereby enhancing spatial learning. By showing that human agents shape not only where we look but how we explore and encode space, this study contributes to a growing understanding of how social meaning guides attention and supports navigation.

**Author summary:** When people explore a new environment, such as an unfamiliar city, they rely on what they see to understand and remember the space. Traditionally, research has focused on stable features like buildings or landmarks. However, real-world environments also include people, whose presence can shape how we explore and learn. In this study, participants explored a virtual city while their eye movements were tracked. Some human figures matched their surroundings, while others appeared out of place. We found that people looked longer at those unexpected figures and that their gaze patterns became more flexible and varied afterward. This broader visual exploration helped them remember the layout of the city more accurately. Our results suggest that human presence, especially when it disrupts expectations, can promote more effective learning by encouraging more complex visual engagement. These findings can provide insight into how strategically placed social cues enhance attention and memory, and more importantly, how they influence our patterns of visual exploration.

## Introduction

Spatial cognition has traditionally been studied through static environmental cues, such as landmarks and architectural layout [1]. These elements provide stable reference points that aid wayfinding and memory encoding. However, real-world spatial cognition extends beyond static structures. Environments contain dynamic elements, including socially relevant cues, that guide spatial understanding. Among these, humans serve as particularly meaningful reference points, as their presence and placement within an environment convey implicit information about its function, affordances, and social relevance [2]. Unlike fixed landmarks, human agents contextualize the environment by embedding scenes with social and semantic meaning. Depending on their relevance to the setting, their presence may also interfere with spatial learning through social cueing [3]. Despite their ubiquity in real-world navigation, the role of fellow humans in shaping spatial knowledge formation remains underexplored.

In virtual environments, recent findings suggest that the mere presence of human agents can enhance spatial awareness and knowledge acquisition [4]. Sánchez-Pacheco et al. systematically investigated the role of human agents in spatial cognition, demonstrating that their presence locally influences virtual space exploration, directs visual attention, and facilitates memory recall [5]. Their work further emphasizes that human agents contribute semantic relevance to an environment by shaping how spatial relationships are perceived. For instance, a construction worker positioned in front of a construction site reinforces the semantic association of that location. In contrast, if the same worker stands in front of a basketball court, the mismatch creates an incongruence that enhances participants’ ability to remember the location. These findings support the importance of the presence of human agents and their spatial and contextual relevance in shaping visual exploration and knowledge acquisition. Yet how this contextual relevance translates into changes in perception and encoding remains to be understood.

A key starting point is vision, which plays a central role in how spatial information is perceived, organized, and remembered. While navigating an environment, individuals direct their visual exploration toward areas of interest before determining goal-oriented body movement [6]. Visual exploration involves a balance between exploitation, where attention is concentrated on task-relevant, familiar elements, and exploration, where fixations shift flexibly to gather new spatial information [7]. Fixations have traditionally been considered indicators of exploitation, as they reflect the selection of known or relevant information. In contrast, transitions between fixations encode the temporal and sequential structure of exploration, capturing how attention is dynamically allocated across a scene [8]. The interplay of visual exploitation and exploration is particularly relevant in understanding how human agents shape visual behavior.

To quantify fixation dynamics, gaze transition entropy (GTE) provides a structured measure of dispersed versus predictable visual exploration [9]. Higher GTE values indicate a more exploratory scanning strategy, while lower values reflect a focused, goal-directed search. Beyond capturing fixation dispersion, GTE has been linked to higher-order cognitive processes, including attentional allocation [10], task difficulty [11], and perceived task proficiency [12]. Social presence has also been shown to influence fixation behavior, either by clustering fixations due to social salience [13] or by modulating them based on contextual relevance [14]. However, it remains unclear how social context, particularly the congruence between agents and their surroundings, shapes variability in visual exploration as captured by GTE.

To address this question, we designed a 1 km^2^ virtual city composed of 236 buildings and 56 human agents. A subset of agents held objects implying specific actions (e.g., a shovel or a donut) and were positioned either in semantically congruent contexts (e.g., a construction worker at a construction site) or incongruent ones (e.g., a construction worker in front of a donut shop). Participants freely explored the city while their gaze behavior was recorded, allowing for the analysis of fixation durations and GTE in response to different contextual conditions.

Here, we investigate how the presence of human agents influences visual exploration and spatial knowledge acquisition in virtual environments. Specifically, we examine whether action-implying agents modulate attention allocation and encoding processes depending on the congruence between agent and environment. We hypothesize that human agents encourage broader exploration by increasing attentional shifts, and that semantic congruence modulates both visual behavior and subsequent spatial learning.

## Materials and methods

### Participants

The dataset analyzed in this study was previously collected and described by Sánchez Pacheco et al. [5]. While the original analysis focused on spatial navigation and memory performance, the present study builds on this dataset by introducing a novel analysis of gaze behavior and fixation dynamics, offering new insight into the visual processes that accompany spatial learning in socially contextualized environments. Briefly, the study consisted of two experiments, for which we initially recruited 70 participants evenly distributed across both experiments. All participants had normal or corrected-to-normal vision. Due to attrition, the final sample comprised 53 participants. Ten participants were unable to complete the study due to illness or scheduling conflicts, three withdrew due to motion sickness, and four were excluded due to incomplete data. The final distribution included 27 participants in Experiment 1 (14 male, 13 female) and 26 in Experiment 2 (11 male, 15 female). In Experiment 1, participants ranged in age from 19 to 46 years (*M* = 23.92, *SD* = 6.72), while in Experiment 2, ages ranged from 19 to 31 years (*M* = 22.29, *SD* = 2.90).

### Experimental design and setup

Participants freely navigated a virtual city while their eye movements were continuously recorded to investigate how contextual agents influence visual exploration and spatial learning. The experimental setup, virtual environment, and procedure followed the methodology outlined in Sánchez-Pacheco et al. [5]. Participants explored a 1 km^2^ virtual city with 236 buildings categorized into residential areas and public spaces, such as restaurants, retail stores, recreational areas, and construction sites (see Fig 1A). Among these, 52 buildings were marked with street art, evenly distributed across residential and public spaces, designating them as task-relevant locations. Four large peripheral buildings functioned as global landmarks, serving as reference points for navigation. This structured environment allowed for controlled manipulation of spatial cues, ensuring visual behavior could be analyzed in response to architectural elements and agent placement.

**Fig 1.**
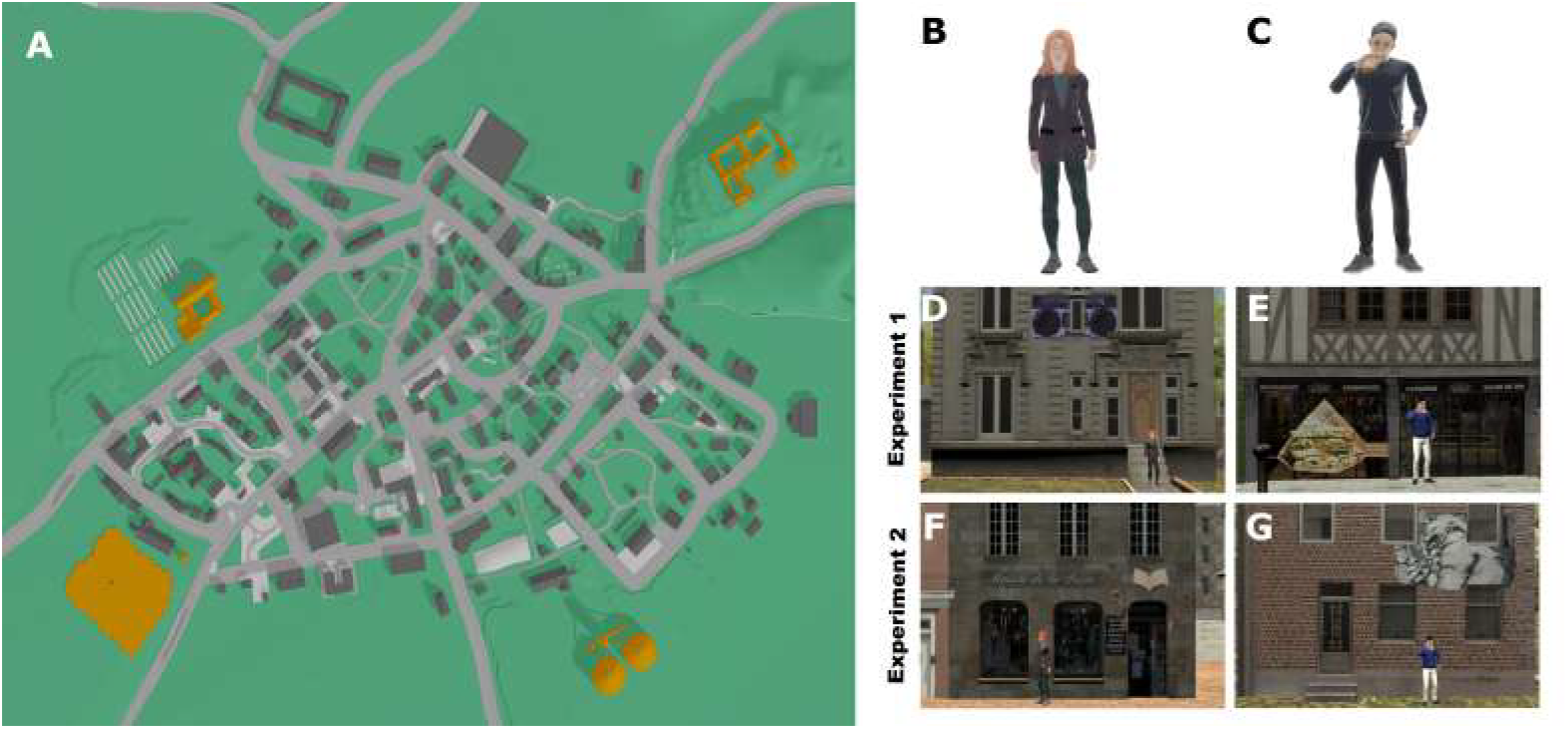
Experimental design and virtual environment. (A) Bird’s-eye view of the 1 km^2^ virtual city, showing buildings and paths. Orange marks indicate the four global landmarks. (B) Example of an acontextual agent. (C) Example of a contextual agent. (D–G) Agent placements across experimental conditions. In Experiment 1, (D) acontextual agents in residential areas, and (E) contextual agents in congruent locations (e.g., a sandwich in front of a sandwich shop). In Experiment 2, (F) acontextual agents in both residential and public spaces, and (G) contextual agents in incongruent settings.

Each participant completed five 30-minute exploration sessions, followed by a final 60-minute test session to assess spatial knowledge. During all sessions, participants were free to explore the city while their eye movements were recorded using a VR-based eye-tracking system. The final session included pointing tasks to evaluate their internal spatial representations. To ensure data quality, eye-tracker calibration was repeated every 10 minutes. This protocol allowed for high-resolution tracking of gaze behavior across repeated exposures, forming the basis for linking visual exploration with spatial learning outcomes.

The exploration and task assessment sessions were conducted with a desktop computer with an Intel^®^ Xeon^®^ W-2133 CPU, 16 GB RAM, and a Nvidia RTX 2080 Ti graphics card. The VR environment was rendered using an HTC Vive Pro Eye head-mounted display (HMD), operating at a refresh rate of 90 Hz with an effective field of view of approximately 110°. We used four SteamVR Base Stations 2.0, an HTC VIVE body tracker 2.0, and Valve Index controllers to monitor participants’ positions within the environment. This combined setup achieved sub-millimeter precision in capturing the head, body, and eye positions, rotation, and orientation.

The key experimental manipulation involved the placement of contextual agents (see Fig 1). Overall, we placed 56 agents throughout the virtual city, 28 contextual and 28 acontextual. In Experiment 1, acontextual agents were placed in neutral residential settings (Fig 1B, D) and contextual agents in congruent locations, where their held objects aligned with the surrounding environment, such as a sandwich in front of a sandwich shop (Fig 1C, E), reinforcing contextual consistency. In Experiment 2, all agents were redistributed: acontextual agents were placed evenly across residential and public spaces, the latter including locations associated with commercial and recreational activities (Fig 1F), while previously congruent agents were relocated to incongruent settings, ensuring they were placed in public or residential areas that did not match their held objects (Fig 1G), thereby disrupting contextual expectations. This manipulation of the virtual environment and agent placement allowed us to investigate how visual behavior systematically adapts to both environmental layout and agent categories, forming the ground for our entropy analyses of human-context-driven spatial knowledge acquisition.

As a final step, all participants completed a sixth session in which spatial knowledge was assessed. In this session, a VR-based pointing task was administered, where participants indicated the direction of target buildings from multiple randomized locations. Performance was measured as the angular error between the indicated and actual locations. The number and distribution of trials differed between experiments (336 and 224 trials, respectively) to accommodate additional tasks in the second experiment (see [5]). This final task provided a quantitative measure of spatial knowledge, where lower angular errors reflected higher accuracy, enabling direct comparisons across participants and experimental conditions.

### Fixation classification and duration accumulation

The virtual environment was constructed with mesh colliders assigned to all objects, enabling tracking of participants’ fixation behavior by projecting 3D eye vectors onto the scene. Fixations and saccades were identified using a velocity-based classification algorithm [15], following the approach outlined in Sánchez-Pacheco et al. [5]. After classifying all gaze data into fixations and saccades, fixation durations were accumulated per object. For each participant, the total fixation time on each task house and agent was computed by summing the cumulative fixation duration across the five exploration sessions. This yielded a measure of total visual engagement per object, enabling analysis of how attention was focused on specific elements in the virtual environment.

### Entropy calculation

Gaze transition entropy was computed to quantify variability in gaze shifts across different environmental elements. Each participant’s transition matrix was generated per session to capture the frequency of fixation transitions between predefined categories. Each fixation was assigned to one of eight visual categories: background elements, buildings, task-relevant residential buildings, task-relevant public buildings, global landmarks, acontextual agents, and contextual agents (i.e., congruent and incongruent). Only forward transitions were considered, ensuring that each fixation was linked exclusively to the subsequent fixation in the sequence (see Fig 2A, B). The resulting matrices were row-normalized to obtain a probability distribution of fixation transitions across categories (Fig 2C).

**Fig 2.**
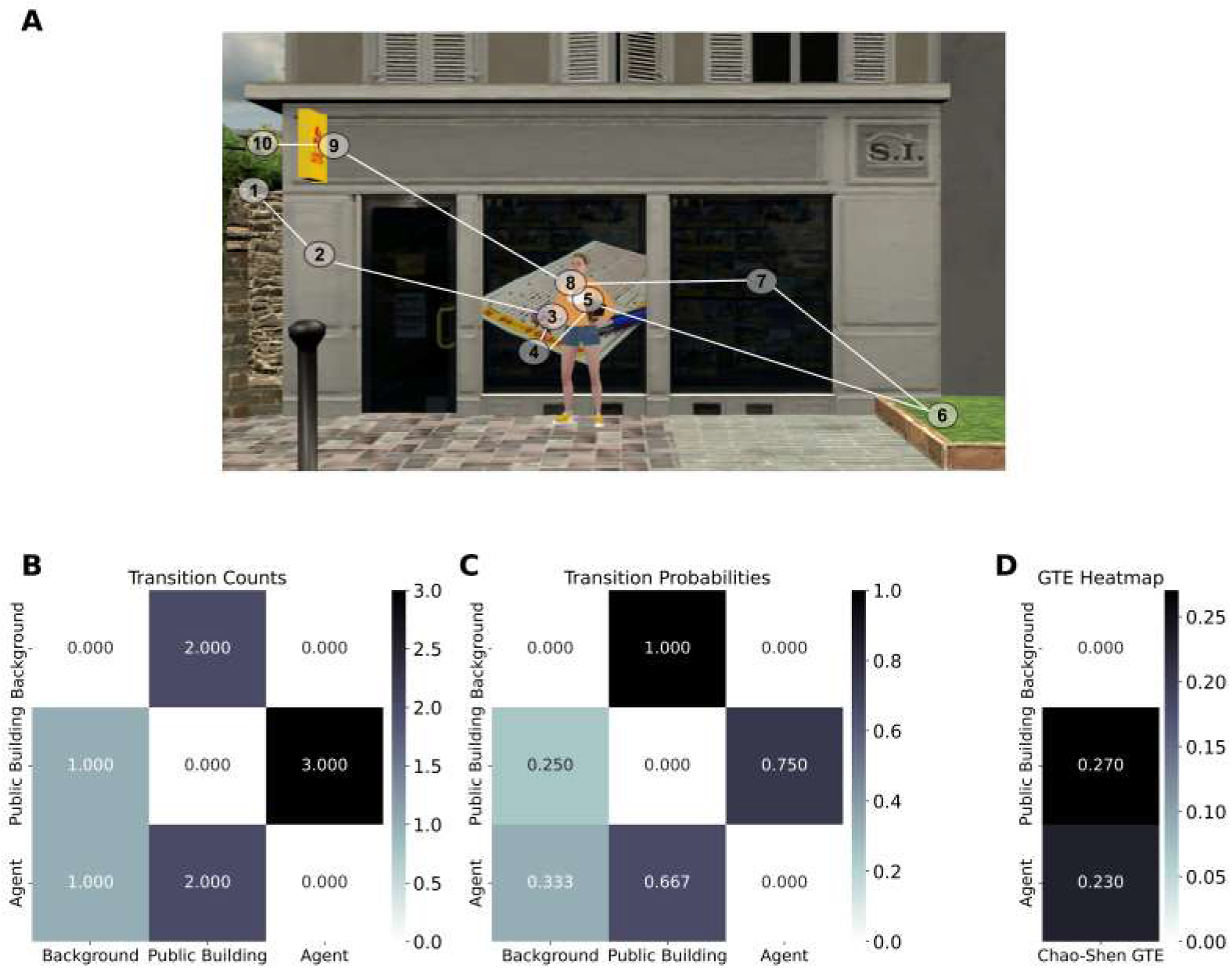
Fixation transitions across visual categories. (A) Sequential gaze transitions between visual categories, with numbered circles indicating fixations. (B) Raw transition counts showing gaze shift frequencies. (C) Row-normalized transition probabilities. (D) Chao-Shen corrected gaze transition entropy (GTE), where higher values reflect more variable scanning.

To quantify fixation variability, we computed entropy from the row-wise transition probabilities of each category using the Chao-Shen estimator [16], which accounts for sample-size bias and rare transitions. This correction specifically adjusts for singleton transitions—categories observed only once—thus improving entropy estimation under sparse sampling conditions. To ensure methodological rigor and comparability with prior eye-tracking research, we followed the implementation approach used by Wilming et al. [17]. Conceptually, this method aligns with missing-mass approximations in neural modeling frameworks, such as those developed by Haslinger et al. [18], which similarly address the influence of unobserved elements on model structure and entropy-based inference. The category-specific corrected entropy was defined as shown in Eq (1):

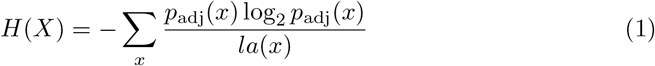

Here, *p*_adj_(*x*) represents the adjusted transition probability for category *x*, and *la*(*x*) is the likelihood of observing that category at least once. The adjustment is based on the estimated sample coverage, computed using the number of singleton transitions *S*_1_ and the total number of transitions *N*, as shown in Eq (2):

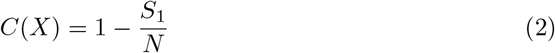

To derive a global measure of gaze entropy across all categories, we computed a weighted average of the category-specific entropies. Each weight corresponds to the stationary distribution of gaze within that category, yielding a global GTE score as defined in Eq (3):

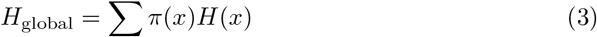

where *π*(*x*) denotes the stationary probability of fixating on category *x*, derived from the normalized transition matrix.

Finally, to allow comparison across participants and conditions, all entropy values were normalized by dividing them by the theoretical maximum entropy given the number of fixation categories *k*. This normalization is shown in Eq (4):

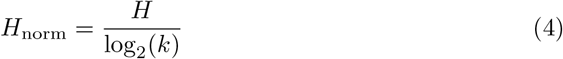

Normalized GTE values ranged from 0 (completely predictable transitions) to 1 (maximally unpredictable, uniform transitions), facilitating cross-subject and cross-session analyses.

### Local entropy: segmented entropy calculation for agent encounters

We introduced a more localized GTE calculation to assess whether gaze variability changed following interactions with human agents. For each detected fixation on a human agent, a 30-second pre-interaction window and a 30-second post-interaction window were defined. Non-overlapping trials were enforced by considering only the first valid fixation on a given collider within 30 seconds. If multiple fixations occurred in rapid succession, only the most recent non-overlapping instance was retained.

Transition matrices were computed separately for each window and normalized to obtain transition probabilities. GTE was then computed using the Chao-Shen correction (see Eq (1)), and values were normalized using Eq (4) to ensure comparability across trials. This method allowed us to systematically assess whether local interactions with agents triggered changes in fixation entropy, indicating shifts in visual exploration strategies.

### Data modeling: statistical analysis

We used Bayesian hierarchical models to analyze fixation behavior and gaze entropy. All models were implemented in the `brms` package (version 2.21.0) in R [19], which interfaces with Stan for Hamiltonian Monte Carlo (HMC) sampling [20]. Each model was run with four chains of 4,000 iterations, including 1,000 to 2,000 warmup iterations.

Model families were chosen to reflect the distributional properties of each outcome variable. Gamma regression with a log link was used for models predicting absolute pointing error and dwell time, both of which are strictly positive and right-skewed. Beta regression with a logit link was used for models predicting normalized GTE, given the bounded nature of entropy values between zero and one. For the Gamma models, weakly informative priors were specified explicitly: Normal priors 𝒩 (0, 1) were assigned to fixed effects, and Cauchy priors Cauchy(0, 2.5) were used for intercepts and group-level standard deviations. Beta models were fit using default priors provided by `brms`. Posterior distributions were summarized using means and 89% highest density intervals (HDIs), following the default convention in `brms`. Model convergence was assessed using the Gelman-Rubin statistic 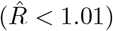 and visual inspection of trace plots.

### Bayesian analysis of dwell time on agents and buildings

To examine how object type and agent congruency influenced dwell time *D*, we fit a Bayesian hierarchical model including fixed effects of object type *ω* and congruency *γ*, as well as random intercepts for participant *ι* and session *σ*, as specified in Eq (5):

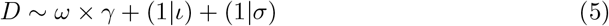

Here, *ω* distinguishes between agents and buildings, while *γ* captures the contextual congruence of the agent (i.e., congruent, incongruent, acontextual). The interaction term allows us to assess whether incongruent agents elicited longer fixations and whether these effects extended to nearby buildings.

### Bayesian time models for entropy dynamics

To model the evolution of fixation entropy *H* across agent encounters, we implemented a beta regression with a logit link, appropriate for bounded outcome variables. The model incorporated encounter index *k* and a smoothing spline *s*(*k*, 5) to account for nonlinear temporal changes, as shown in Eq (6):

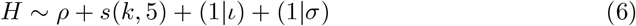

In this model, *ρ* contrasts pre- and post-encounter entropy values, while the spline *s*(*k*, 5) captures smooth transitions across successive interactions. Participant- and session-level random intercepts again account for individual differences.

### Post-encounter entropy model

To isolate the effects of agent context on post-interaction gaze variability, we specified a third model restricted to post-fixation GTE values. This model evaluated whether the type of agent encountered influenced subsequent exploration patterns (Eq (7)):

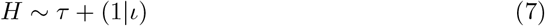

Here, *τ* denotes agent context condition (Congruent, Incongruent, Acontextual), and *ι* is the participant-level random intercept. This model was fit to a filtered dataset containing only post-gaze entropy estimates. Together, this series of models enables a nuanced understanding of how visual exploration is shaped by contextual relevance, temporal dynamics, and individual variation.

## Results

To examine how contextual agents influenced gaze behavior and spatial knowledge, we analyzed fixation durations, gaze transition entropy (GTE), and pointing accuracy using Bayesian hierarchical models. These analyses were conducted on gaze data recorded across five exploration sessions and spatial memory assessments from a final test session. Fixation metrics were computed at both global and local levels, with separate models addressing cumulative dwell time, entropy dynamics over time, and post-encounter entropy changes. All models included participant- and session-level random effects to account for individual variability. Below, we present the results in three parts: (1) fixation duration and engagement across agent conditions, (2) temporal modeling of entropy dynamics, and (3) post-interaction changes in visual exploration.

### Fixation duration and engagement across agent conditions

To examine whether agent congruency influenced attentional engagement, we analyzed total dwell time and fixation counts on agents and associated buildings across conditions (Fig 3). We hypothesized that incongruent agents would elicit the greatest attention, followed by congruent and acontextual agents, and that these effects might extend to nearby buildings. Visual engagement was assessed through descriptive statistics and a Bayesian hierarchical regression model.

**Fig 3.**
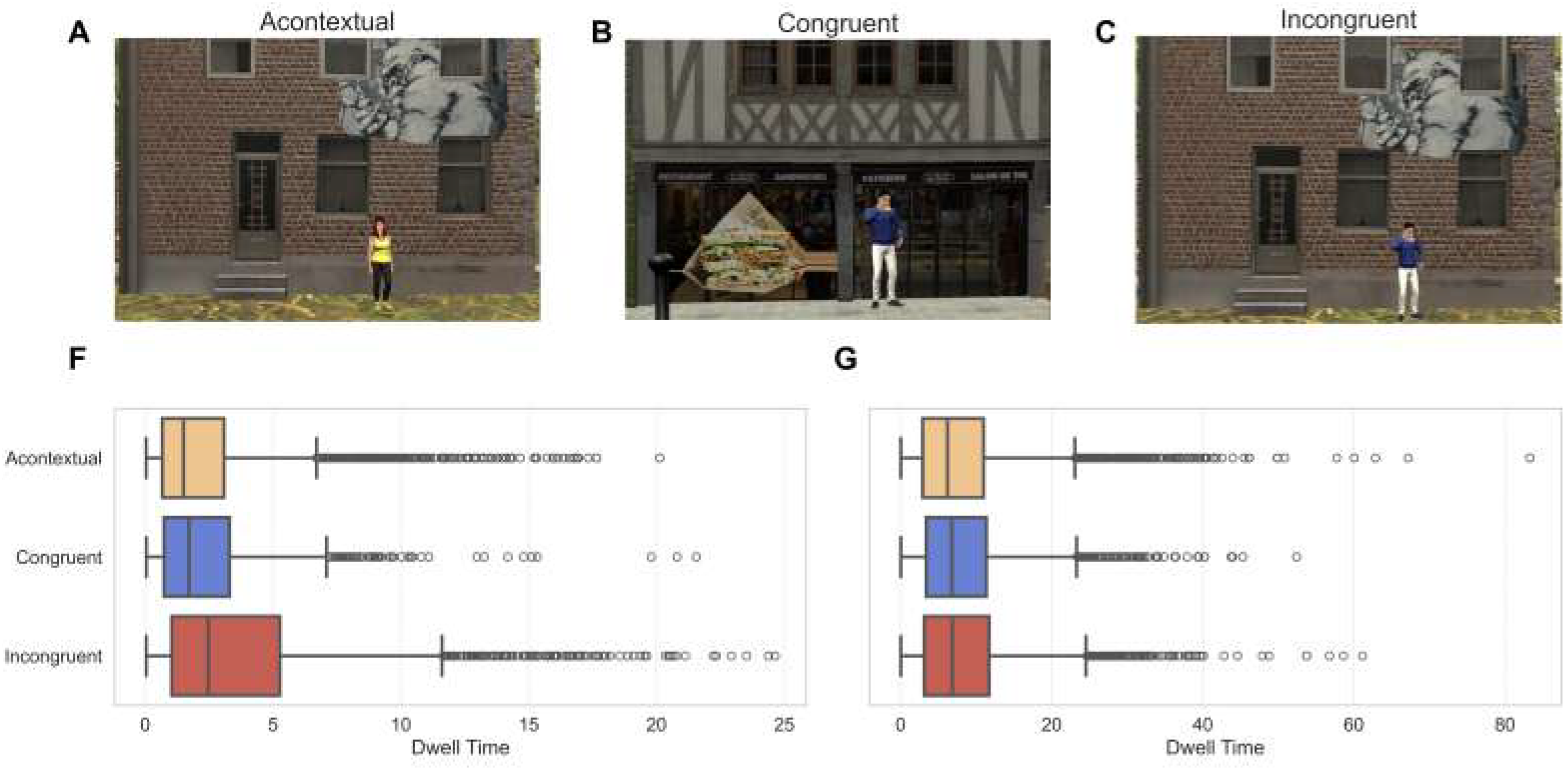
Agent types and dwell time distributions. (A–C) The three agent conditions: (A) Acontextual, (B) Congruent, and (C) Incongruent. In (B), the agent’s object matches the environment; in (C), it does not. (D–E) Boxplots of dwell time for (D) agent fixations and (E) building fixations. (F–G) Model-derived estimates with 89% HDIs.

Dwell time is a common proxy for attentional allocation, with longer durations reflecting greater scrutiny or interest [21]. Incongruent agents attracted significantly longer fixations (M = 3.83 s, SD = 3.90) than both acontextual (M = 2.36 s, SD = 2.57) and congruent agents (M = 2.26 s, SD = 2.50) (Fig 3D,F). Similarly, fixation counts were highest for incongruent agents (M = 7.90, SD = 10.52), compared to acontextual (M = 5.19, SD = 6.75) and congruent agents (M = 5.09, SD = 7.53). These patterns suggest increased visual engagement with incongruent agents, likely due to their violation of contextual expectations.

We next assessed whether agent congruency influenced attention toward nearby buildings. Participants spent a comparable amount of time fixating on buildings near congruent (M = 8.20 s, SD = 6.54) and incongruent agents (M = 8.50 s, SD = 7.40), while slightly less time was spent near acontextual agents (M = 8.00 s, SD = 7.13) (Fig 3E,G). Fixation counts followed a similar trend: 14.90 (SD = 14.18) for congruent, 15.22 (SD = 15.54) for incongruent, and 14.75 (SD = 14.81) for acontextual conditions. These small differences suggest that while incongruent agents attracted greater direct attention, spillover effects on nearby buildings were less pronounced.

To formally assess the effect of agent congruency and object type on visual engagement, we fit a Bayesian hierarchical model using a Gamma distribution with a log link. The model included fixed effects for object type (agent vs. building), congruency, and their interaction, as well as random intercepts for participant (*n* = 57) and session number (*n* = 5). The shape parameter confirmed a right-skewed distribution of dwell times (*β* = 1.43, 89% HDI: [1.40, 1.45]).

Agents were associated with significantly shorter dwell times overall (*β* = *−*1.16, 89% HDI: [−1.18, −1.13]), reflecting a 68.3% reduction compared to buildings. Incongruency had a small but positive effect on overall dwell time (*β* =*−* 0.07, 89% HDI: [0.02, 0.12]), corresponding to a 7.25% increase. Importantly, this effect was moderated by object type (*β* = −0.40, 89% HDI: [−0.47, −0.34]), such that incongruency increased dwell time for agents but decreased it for buildings. Specifically, incongruent agents received 30% longer fixations, whereas nearby buildings were fixated on 13% less.

These results reveal a directional asymmetry in attention allocation: participants engaged more strongly with socially incongruent agents but showed reduced fixation on their surrounding context. This pattern suggests that violations of social expectations capture attention more effectively than mismatches between an object and its environment.

### Gaze transition entropy across visual categories and congruency

To assess whether agent congruency influenced the fluidity of visual exploration, we examined GTE across visual categories using Chao-Shen corrected entropy estimates (Fig 4). We hypothesized that incongruent agents would disrupt attentional flow, leading to greater variability in gaze transitions.

**Fig 4.**
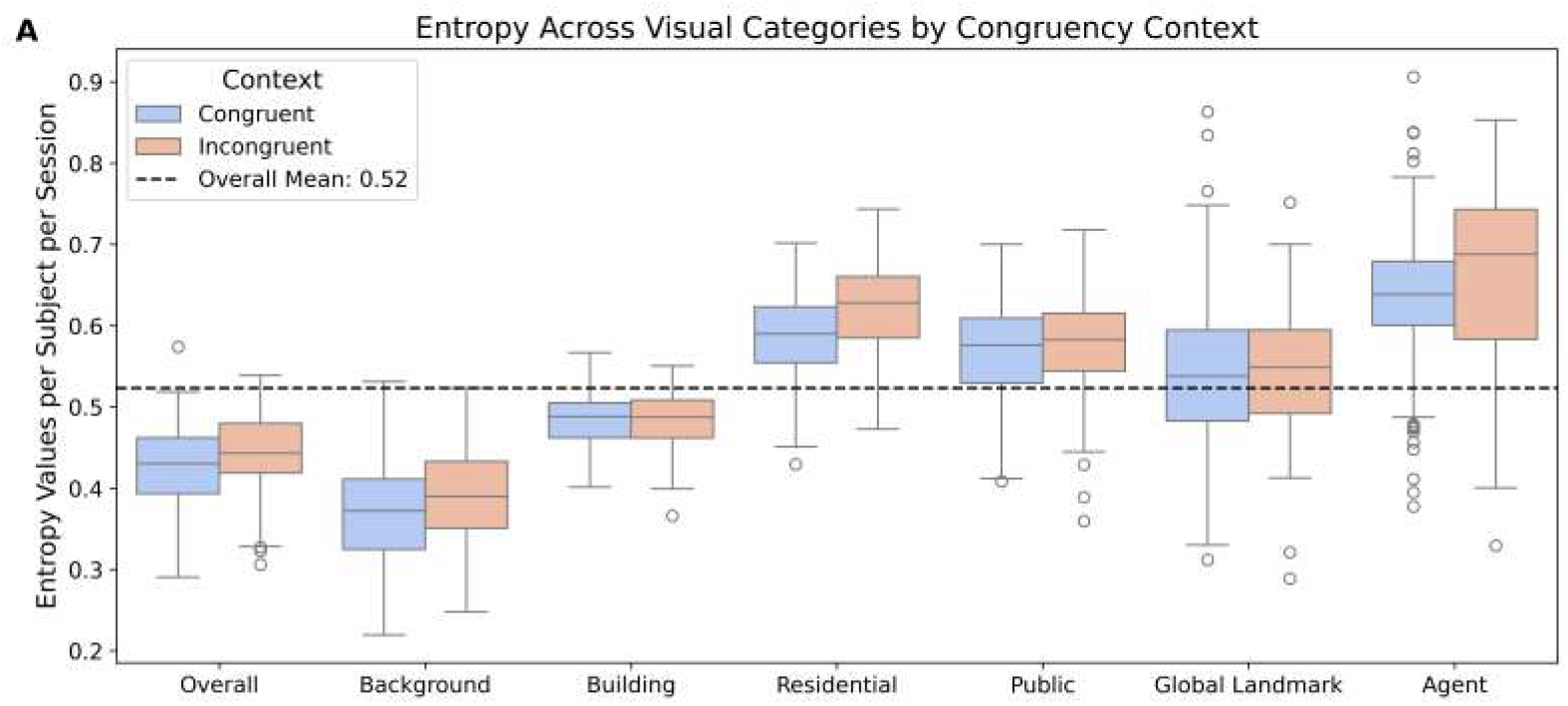
Categorical gaze transition entropy across visual categories and contexts. (A) Boxplots of Chao-Shen corrected gaze transition entropy (GTE) per subject per session, split by visual category and condition. Higher values indicate more unpredictable gaze transitions. Dashed line indicates overall mean.

On average, overall entropy was higher in the incongruent condition (*M* = 0.44, *SD* = 0.05) than in the congruent condition (*M* = 0.42, *SD* = 0.05), suggesting that incongruency modestly increased the dispersion of gaze patterns. This effect was most pronounced when focusing on agents. GTE for agents rose from *M* = 0.63, *SD* = 0.09 in the congruent condition to *M* = 0.67, *SD* = 0.11 in the incongruent condition, reflecting more exploratory and less predictable transitions following encounters with incongruent agents.

In contrast, entropy estimates for other categories were largely stable across conditions. Buildings showed no meaningful change (*M* = 0.48, *SD* = 0.03 congruent; *M* = 0.48, *SD* = 0.04 incongruent), and global landmarks also remained constant (*M* = 0.54, *SD* = 0.10 vs. *M* = 0.54, *SD* = 0.07).

These findings indicate that incongruent agents uniquely disrupt gaze dynamics, increasing transition entropy and prompting more variable patterns of visual exploration. Other visual elements, including buildings and landmarks, were comparatively unaffected by agent congruency, suggesting a targeted influence of social-contextual anomalies on attentional behavior.

### Exploration–exploitation trade-off: The effect of agent congruency

To investigate how agent congruency modulated attentional trade-offs between exploration and exploitation, we analyzed the relationship between dwell time and gaze transition entropy using kernel density estimation (KDE) and correlation analysis (Fig 5). We expected incongruent agents to elicit prolonged and less predictable gaze behavior—indicative of greater exploration.

**Fig 5.**
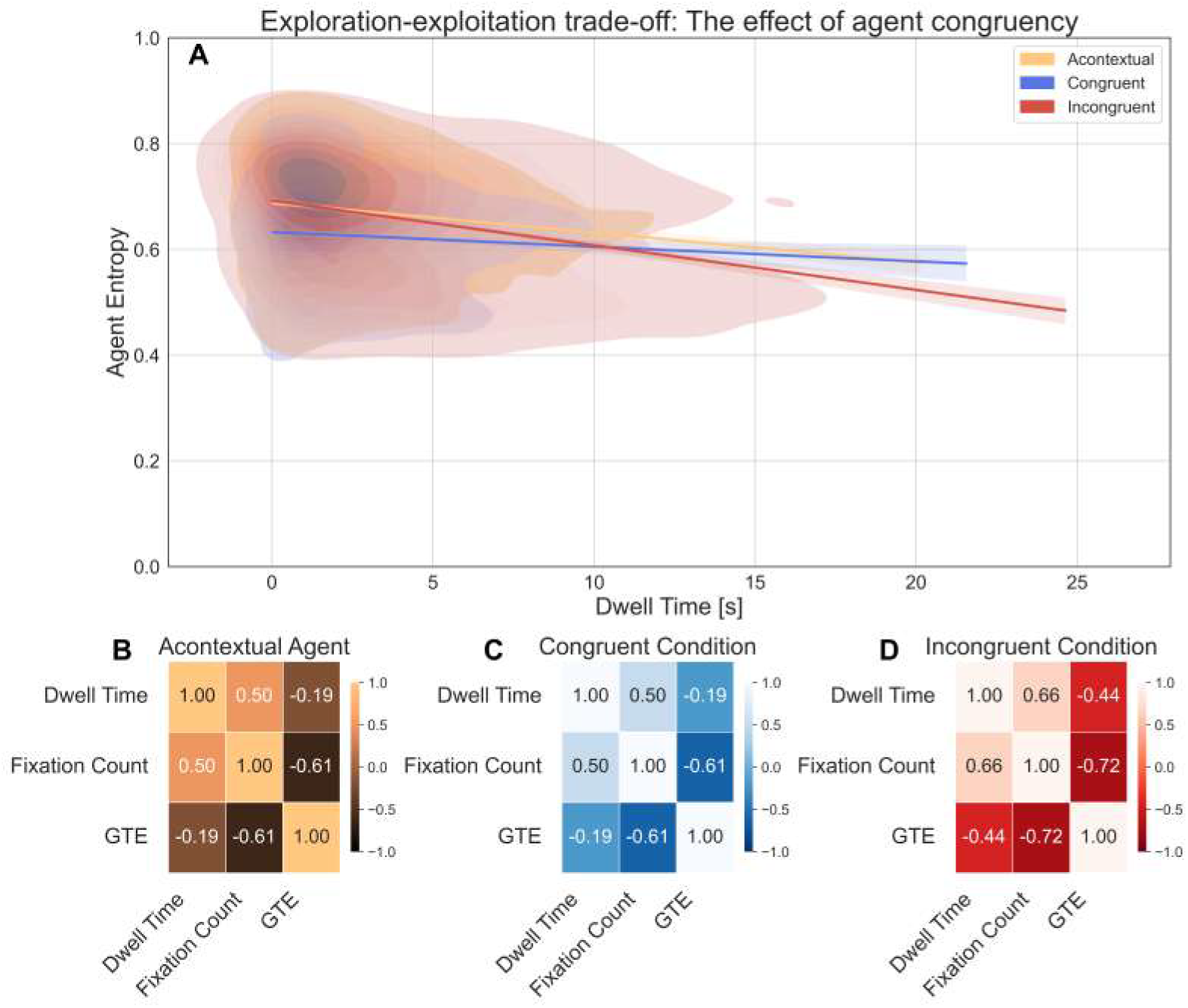
Relationship between dwell time, gaze transition entropy, and fixation count for agent categories. (A) Kernel density estimation (KDE) plot of dwell time vs. GTE for (yellow) acontextual, (blue) congruent, and (red) incongruent agents. Regression lines show trends per condition. (B–D) Correlation matrices of dwell time, fixation count, and GTE.

In Fig 5A, KDE analysis showed that incongruent agents had the highest mode of dwell time (1.17 s) and centroid (4.19 s), suggesting both common and average fixations were longer compared to congruent (mode = 0.80 s, centroid = 3.21 s) and acontextual agents (mode = 0.76 s, centroid = 2.96 s). Additionally, incongruent agents showed the widest dwell time distribution (FWHM = 3.82 s), reflecting greater variability than congruent (2.69 s) and acontextual agents (2.31 s).

Entropy patterns mirrored this trend: incongruent agents showed the highest mode (0.735) and centroid (0.661), indicating more dispersed gaze behavior. Congruent agents exhibited the lowest entropy (mode = 0.637, centroid = 0.626) with the narrowest spread (FWHM = 0.094), suggesting fixations on them were more structured and predictable. These results suggest that incongruence increased both the duration and variability of gaze, reflecting a shift toward more exploratory viewing strategies.

To assess the relationship between visual exploitation and exploration, we computed correlation matrices between dwell time, fixation count, and GTE (Fig 5B–D). For acontextual agents, fixation count and dwell time were moderately correlated (*r* = *−*0.50), while both were negatively associated with GTE (*r* = *−*0.61; *r* = −0.19), suggesting that greater local attention tended to co-occur with more structured gaze patterns.

In the congruent condition, fixation count and dwell time were similarly correlated (*r* =*−* 0.50), and both showed weaker negative correlations with GTE (*r* = *−*0.61; *r* = *−*0.19), indicating a similar but less pronounced trade-off between attention and entropy.

In the incongruent condition, these trade-offs were most pronounced. Fixation count and dwell time were strongly correlated (*r* = *−*0.66), while their correlations with GTE were sharply negative (*r* = *−*0.72; *r* = −0.44). These findings suggest that incongruent agents elicited a robust attentional capture effect, where prolonged fixations were tightly coupled with reduced gaze variability—marking a sharper shift toward exploitation. Together, these results highlight how agent congruency modulates attentional dynamics: incongruence enhances dwell time while simultaneously suppressing exploratory gaze, producing the strongest exploration–exploitation trade-off across conditions.

In the congruent condition, fixation count and dwell time were also positively related (*r*(128) = *−*0.50, *p <* .001), but both were negatively correlated with GTE (*r* =*−* 0.19, *p* = .029; *r* = *−*0.61, *p <* .001), indicating that increased attention corresponded to more constrained gaze transitions—consistent with exploitation.

In the incongruent condition, these trade-offs were most pronounced. Dwell time and fixation count correlated strongly (*r*(145) = *−*0.66, *p <* .001), while their relationships with entropy were sharply negative (*r* = *−*0.44, *p <* .001; *r* = −0.72, *p <* .001). These findings suggest that incongruent agents elicited a robust attentional capture effect, where prolonged fixations were tightly coupled with reduced gaze variability—marking a sharp shift toward exploitation. Together, these results highlight how agent congruency modulates attentional dynamics: incongruence enhances dwell time while simultaneously suppressing exploratory gaze, producing the strongest exploration–exploitation trade-off across conditions.

### Time analysis of GTE: Agent effects on entropy

To assess whether agent encounters dynamically influenced visual exploration, we analyzed gaze transition entropy (GTE) before and after fixating on an agent. A Bayesian hierarchical regression model incorporating a smoothing spline for event index and a multilevel structure for individual and session variation was used to examine entropy trajectories over time.

As shown in Fig 6A, post-fixation entropy significantly increased on the logit scale (*β* = 0.37, 95% CI = [0.35, 0.38]). Back-transforming using the model’s intercept (*α* = 0.75) suggests entropy rose from approximately 0.68 (pre-fixation) to 0.75 (post-fixation), a relative increase of 10.9%. This indicates that gaze transitions became more dispersed and less predictable after fixating on agents. The smoothing spline for event index showed a non-significant effect (*β* = 0.35, 95% CI = [–0.03, 0.79]), suggesting no systematic drift over successive encounters.

**Fig 6.**
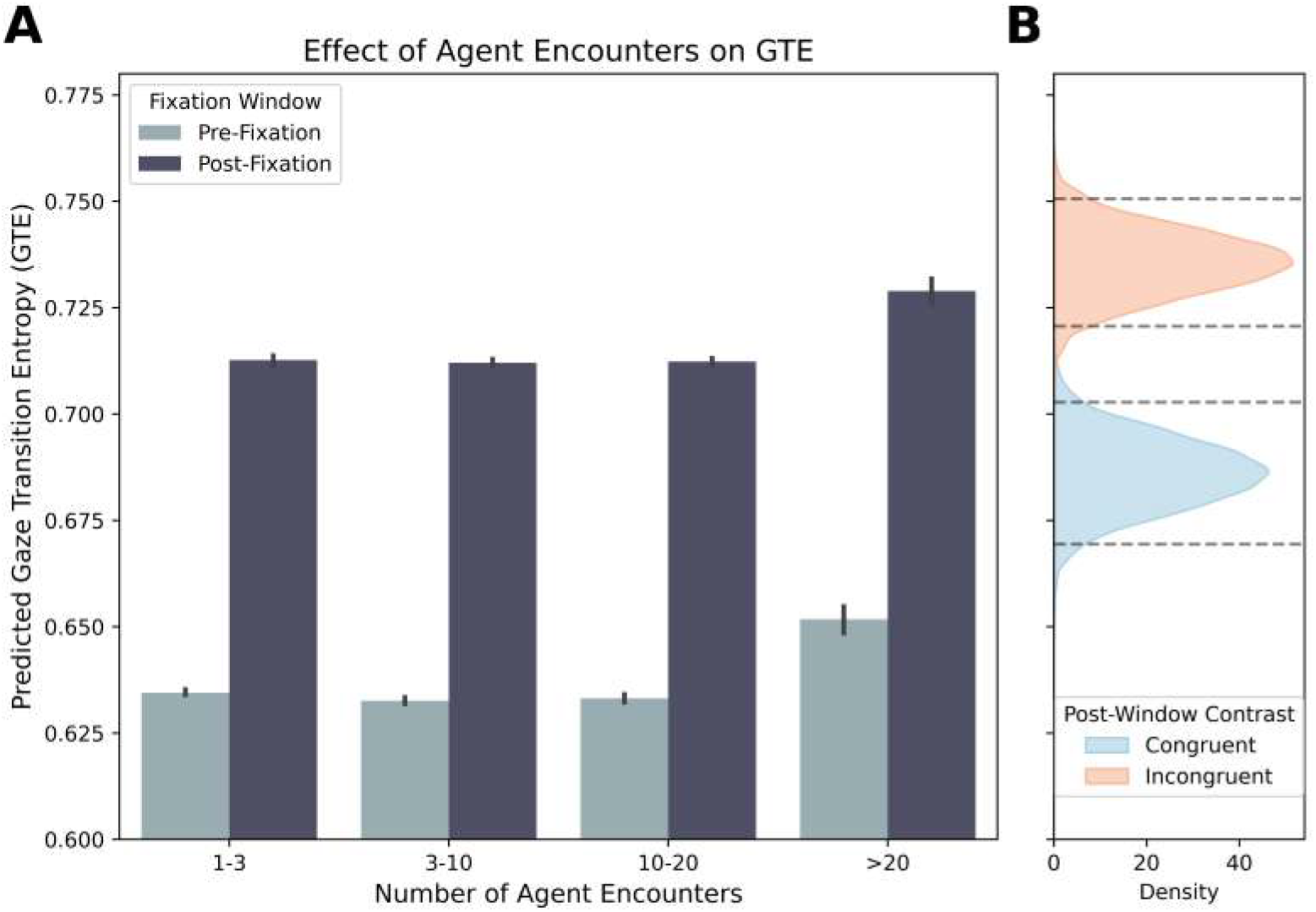
Effect of agent encounters on gaze transition entropy. (A) Model-based predictions of GTE before and after agent fixations. Shaded ribbons show uncertainty. (B) Posterior distributions from post-fixation model comparing agent types. Dashed lines mark 95% CIs.

To ensure effects reflected genuine changes rather than temporal drift, pre- and post-fixation windows were defined as non-overlapping 30-second segments. Modest variability in participant- and session-level intercepts (*SD*_ID_ = 0.23; *SD*_Session_ = 0.12) confirmed baseline stability. A Bayes factor of *BF*_10_ *≈*3.39 *×* 10^93^ strongly favored the full model, confirming a real increase in entropy after agent encounters.

To isolate the influence of agent type, we fit a second Bayesian beta regression model to post-fixation GTE, using agent type as a categorical predictor (Fig 6B). The model intercept for acontextual agents was *β* = 0.94 (95% CI = [0.89, 1.00]), corresponding to a mean GTE of 0.72. Compared to this baseline, congruent agents significantly decreased post-fixation entropy (*β* = −0.16, 95% CI = [–0.22, –0.11]), yielding a back-transformed mean of 0.68. In contrast, incongruent agents increased post-fixation entropy (*β* = *−*0.08, 95% CI = [0.03, 0.13]), with a mean of 0.74. These results show that congruent agents lead to more focused gaze patterns, while incongruent agents trigger more exploratory viewing behavior. Together, these findings reveal that agent congruency significantly modulates post-fixation gaze behavior, with incongruent cues promoting exploration and congruent cues reinforcing structured, goal-directed scanning.

### Performance prediction using entropy and dwell times

To evaluate how gaze dynamics contribute to spatial learning, we implemented a Bayesian hierarchical gamma regression model to predict absolute pointing error based on post-encounter gaze entropy, dwelling time, and agent context. This approach builds on the notion that effective spatial learning may depend on a balance between exploratory and exploitative gaze behavior: dispersed fixations facilitate the integration of spatial relationships, while focused attention supports precise encoding. Random intercepts were included for participants and pointing task starting locations to account for individual and environmental variability. Participant-level variance exceeded that of location-level effects (*SD*_Subject_ = 0.34, 95% CI = [0.28, 0.41]; *SD*_Location_ = 0.19, 95% CI = [0.14, 0.25]), indicating that individual differences played a greater role in performance outcomes.

Turning to the fixed effects (Fig 7), the model revealed that increased gaze transition entropy was significantly associated with lower pointing error (*β* = −0.31, 95% CI = [–0.48, –0.13]). This corresponds to a 27% reduction in error (exp(−0.31) ≈ 0.73), suggesting that more dispersed gaze patterns foster flexible spatial representations.

**Fig 7.**
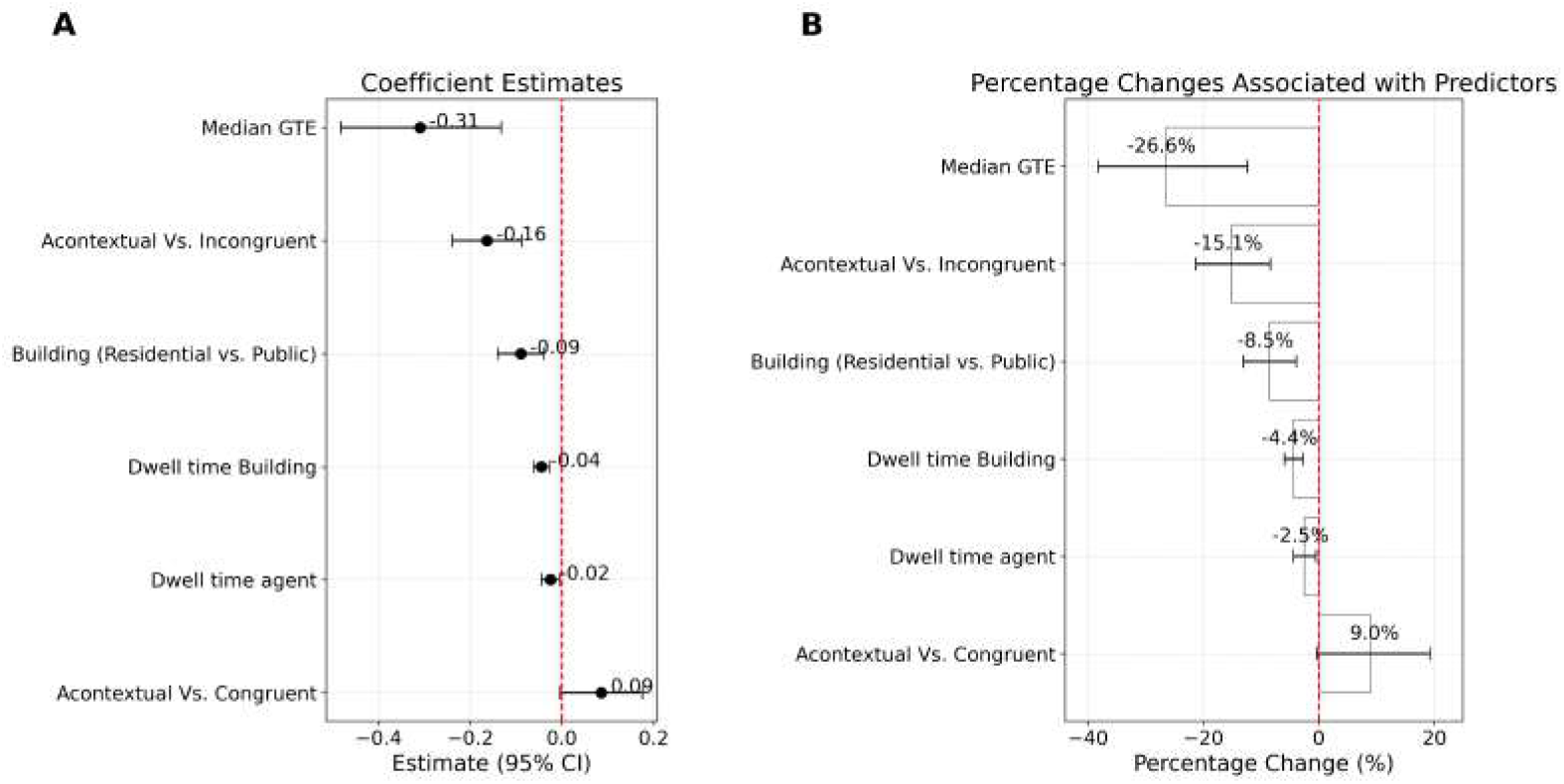
Predicting spatial recall from gaze behavior. (A) Regression surface showing relationship between GTE and pointing error, conditioned on agent dwell time. (B) Posterior estimates of fixed effects including gaze entropy, dwelling times, and contextual contrasts. 95% credible intervals shown.

Fixation duration also contributed to performance. Dwelling longer on buildings reduced pointing error (*β* = –0.04, 95% CI = [–0.06, –0.03]), reflecting a 4% improvement per unit increase. Dwelling on agents showed a similar, albeit smaller, effect (*β* = –0.02, 95% CI = [–0.04, –0.01]), translating to a 2% reduction. These findings support the value of sustained fixations in encoding task-relevant cues, though their effect was modest compared to entropy.

Contextual features also influenced recall accuracy. Participants performed better in public environments relative to residential ones (*β* = –0.09, 95% CI = [–0.14, –0.04]), corresponding to a 9% error reduction (exp(– 0.09) ≈0.91).

Finally, we examined the impact of agent congruency. Relative to the acontextual baseline, the congruent condition yielded a small, non-significant positive effect (*β* = −0.09, 95% CI = [–0.00, 0.18]), while the incongruent condition significantly reduced pointing error (*β* = −0.16, 95% CI = [–0.24, –0.09]), amounting to a 15% improvement (exp(−0.16) ≈0.85). This suggests that incongruent agents—despite violating expectations—may enhance memory through increased gaze dispersion or salience.

In summary, these results highlight gaze entropy as a robust predictor of spatial recall accuracy. While prolonged fixations on agents and buildings also improved performance, entropy-driven exploration exerted the strongest effect. Furthermore, agent incongruency and public context facilitated better recall, reinforcing the importance of both attentional dynamics and environmental features in shaping spatial memory.

## Discussion

This study investigated how human agents influenced visual exploration and learning within a controlled virtual reality city, focusing on how agent-context congruency modulates gaze behavior. The results revealed that agents significantly shaped visual exploration patterns, especially when incongruent with their surroundings. Incongruent agents attracted longer dwell times and increased gaze transitional entropy, promoting more dispersed and less predictable exploration patterns. This suggests that contextual mismatches between agents and their environment trigger attentional shifts beyond the immediate fixation, leading to more distributed scanning behavior. Crucially, gaze transitional entropy, not just experimentally manipulated factors or dwell time, emerged as the most robust predictor of spatial learning performance. Higher entropy during exploration was associated with more accurate spatial recall. In sum, the findings highlight the role of social-contextual information in shaping gaze behavior during spatial learning, demonstrating that incongruent agents enhance visual exploration and, consequently, support the acquisition of spatial knowledge.

While these findings offer new insights into how social agents modulate visual behavior and spatial learning, several limitations should be acknowledged. First, although participants were free to navigate the virtual environment, all objects and agents were non-animated, and the environment itself, while realistic, remained a simplified model of an urban space. This may limit the generalizability of the results to real-world navigation, where movement and environmental dynamics are known to influence attentional allocation [22,23]. However, introducing motion selectively was not feasible, as only contextual agents were equipped with objects or implied actions. Adding animation would have introduced an additional confound, undermining the internal validity of the congruency manipulation. To avoid this, a non-animated environment was employed, ensuring that observed differences in gaze behavior could be attributed specifically to agent-context congruency. Future research could build on these findings by incorporating more dynamic and complex environments, allowing a deeper understanding of how social and contextual factors interact with gaze behavior under motion.

Second, while key visual properties such as agent size, gender, and skin color were controlled, and the distribution of graffiti was balanced across conditions, not all low-level visual features were fully standardized. Agent posture and object properties were left unconstrained to preserve ecological validity and the effectiveness of the congruency manipulation. Although gaze allocation is sensitive to perceptual features such as color or contrast [24], evidence from both experimental and computational studies indicates that gaze behavior in naturalistic tasks is primarily shaped by behavioral goals and semantic relevance [25, 26]. This is especially relevant for entropy-based metrics, as gaze entropy reflects the interaction of bottom-up and top-down processes, with higher entropy often associated with cognitive control and goal-directed exploration [11, 12]. Therefore, while low-level salience may have influenced gaze behavior to some extent, the observed modulation of gaze transition entropy by agent congruency likely reflects meaningful, task-driven exploration dynamics supporting spatial learning.

Agent-context congruency shaped gaze behavior along four distinct empirical dimensions. First, incongruent agents attracted longer fixation durations than both congruent and acontextual agents, suggesting that contextual violations elicited increased local attentional engagement, consistent with previous research showing that semantic inconsistencies prolong fixation times [27]. Second, incongruent agents were associated with higher gaze transition entropy, reflecting a shift from localized and predictable exploration to more distributed and flexible scanning patterns, aligning with findings that expectancy violations induce broader visual exploration [28]. Third, incongruent agents intensified the exploration-exploitation trade-off. Specifically, the negative relationship between dwell time and entropy was stronger for incongruent agents, indicating that prolonged fixations on these agents were systematically followed by increased gaze dispersion across the environment. Finally, this reorganization of gaze patterns had a direct and robust effect on spatial learning. Gaze transition entropy emerged as the strongest predictor of spatial recall performance, surpassing the predictive value of dwell time alone. In sum, agent-context congruency systematically shaped participants’ gaze behavior, accounting for their performance in the spatial memory task.

By violating contextual expectations, incongruent agents not only prolonged fixations on themselves but also promoted a transition from localized exploitation to more distributed and flexible exploration, as reflected in increased gaze transition entropy. This is consistent with the idea that gaze behavior is sensitive not only to the physical properties of stimuli but to their social and contextual significance [29, 30]. In this study, social cues in the environment did not merely anchor visual attention but regulated its distribution across space, redirecting participants’ exploration. While previous studies have demonstrated that agents can serve as attentional anchors during navigation, directing gaze to semantically relevant locations or supporting perspective-taking [4, 5], the present findings indicate that agents can induce redistribution of visual attention when they violate environmental expectations. This aligns with evidence that expectancy violations lead to extended search behavior and attentional shifts [31], particularly when embedded in socially meaningful contexts. Taken together, these findings suggest that agents function not only as attentional landmarks but also as modulators of exploratory behavior, shaping the dynamics of spatial information sampling through their interaction with contextual expectations.

Although all agents were visually identical across conditions—holding the same objects and adopting the same postures—participants systematically adjusted their gaze behavior depending on the agent’s congruency with the environment. This indicates that gaze behavior was not driven solely by visual properties of the agents themselves, but by their contextual embedding within the scene. Rather than being processed as isolated objects, agents appeared to function as relational elements whose meaning emerged from the broader spatial structure. This interpretation is consistent with classical and contemporary accounts of scene processing, which emphasize that comprehension involves not only object recognition but also the integration of functional and contextual relationships between elements [32, 33]. Supporting findings from research on semantic scene understanding [34] and schema-guided attention [35] further suggest that participants interpreted agents in relation to their surrounding context. Congruent agents were smoothly integrated into spatial representations, whereas incongruent agents disrupted these representations and triggered increased exploratory gaze behavior, potentially reflecting attempts to resolve or reinterpret unexpected contextual input. This aligns with prior evidence that semantic violations elicit attentional shifts and enhance memory encoding [36], likely by enriching internal representations. These findings suggest that agent congruency influenced not only where participants looked, but how visual input was organized in relation to the broader scene.

These findings contribute to the growing discussion on the role of humans as environmental facilitators during spatial learning [37, 38]. Previous work has found that agents provide semantic structure, guide attention, and support memory formation during navigation. Specifically, humans in an environment do not merely act as static referential cues, but their presence influences the dynamics of spatial behavior and decision-making [5]. For example, socially relevant agents can serve as meaningful landmarks, shape route choices, support spatial perspective-taking, and facilitate memory consolidation [4, 39]. Evidence from both real and virtual navigation contexts has shown that social relevance enhances gaze allocation [40, 41] and improves spatial perspective-taking compared to purely directional or non-social cues [42]. These results extend these findings by showing that agents influence the selection of fixation targets and the very structure of exploration. This distinction is critical, as it suggests that the benefits of social agents during navigation do not stem solely from their semantic content but from the structural reorganization they induce in visual behavior during spatial exploration.

## Conclusion

In conclusion, spatial cognition is not solely driven by geometry or task demands but is fundamentally shaped by the social meanings embedded in our surroundings. In our study, the presence of incongruent human agents increased the visual entropy of participants’ gaze patterns, encouraging more diverse exploration and promoting deeper engagement with the spatial layout. These findings suggest that even weak human cues, those not directly tied to navigational goals, can scaffold learning. In naturalistic environments, such cues are rarely stable or explicit; they are layered, fleeting, and shaped by the presence and behavior of others. Understanding how these ephemeral signals influence attention and memory is essential for building ecologically valid models of spatial learning. By showing that subtle human incongruence can enhance spatial knowledge through its effects on visual exploration, this study contributes to a growing body of work redefining how context, attention, and cognition interact in real-world-like settings.

## Acknowledgments

The authors sincerely thank all those who supported this project. Special appreciation goes to Nora Maleki and Linus Tiemann for their contributions to the development of the VR city and the Pointing Tasks used in the experiments. They also thank Philipp Spaniol for his 3D artwork and for implementing differential level-of-detail loading for the agents within the scene. Lastly, we thank Melissa Sarria-Mosquera and Kaya Gärtner for their support in data collection.

